# A Genome-Wide Association Study Identifies SNP Markers for Virulence in *Magnaporthe oryzae* Isolates from Sub-Saharan Africa

**DOI:** 10.1101/418509

**Authors:** Veena Devi Ganeshan, Stephen O. Opiyo, Samuel K. Mutiga, Felix Rotich, David M. Thuranira, Vincent M. Were, Ibrahima OuéDraogo, BO Zhou, Darren M. Soanes, James C. Correll, Guo-Liang Wang, Nicholas J. Talbot, Thomas K. Mitchell

**Affiliations:** Department of Plant Pathology, The Ohio State University-Columbus, USA.; Department of Plant Pathology, University of Arkansas-Fayetteville, USA.; Biosciences Department, Exeter University, UK.; Institute of Environment and Agricultural Research, Bobo-Dioulasso, Burkina Faso.; International Rice Research Institute (IRRI), Philippines.

## Abstract

The fungal phytopathogen *Magnaporthe oryzae* causes blast disease in cereals such as rice and finger millet worldwide. In this study, we assessed genetic diversity of 160 isolates from nine sub-Saharan Africa (SSA) and other principal rice producing countries and conducted a genome-wide association study (GWAS) to identify the genomic regions associated with virulence of *M. oryzae*. GBS of isolates provided a large and high-quality 617K single nucleotide polymorphism (SNP) dataset. Disease ratings for each isolate was obtained by inoculating them onto differential lines and locally-adapted rice cultivars. Genome-wide association studies were conducted using the GBS dataset and sixteen disease rating datasets. Principal Component Analysis (PCA) was used an alternative to population structure analysis for studying population stratification from genotypic data. A significant association between disease phenotype and 528 SNPs was observed in six GWA analyses. Homology of sequences encompassing the significant SNPs was determined to predict gene identities and functions. Seventeen genes recurred in six GWA analyses, suggesting a strong association with virulence. Here, the putative genes/genomic regions associated with the significant SNPs are presented.

## INTRODUCTION

Rice is of increasing importance as a staple food crop in sub-Saharan Africa (SSA), particularly due to the rapidly growing urban population. (Amberwaves; Nigatu *et al.,* 2017 USDA, 2017). Currently, 80% of rice production in SSA takes place in eight countries: Nigeria, Madagascar, Mali, Guinea, Côte d’Ivoire, Tanzania, Sierra Leone and Senegal with 20% from the rest of SSA. Nigeria and Madagascar alone account for a third of SSA rice production. It is estimated that rice consumption in SSA will grow from the current 27-28 million tonnes to 36 million tonnes by the year 2026 (Amberwaves; Nigatu *et al.,* 2017 USDA, 2017). Considering the predictions of rice consumption required to attain self-sufficiency, SSA will need to increase production by at least 10% per year for the next 10 years, or succumb to international rice trade with large imports impacting the economy of already financially fragile countries (Amberwaves; Nigatu *et al.,* 2017 USDA, 2017). Because of the increase in demand, several strategies are underway to increase rice production, which include increased usage of fertilizers, planting high-yielding cultivars and increasing the area of rice production (Balasubramanian *et al.,* 2007). In spite of efforts to increase production, biotic constraints such as damage by fungal pathogens, remain a major challenge towards achieving this food security goal. Recent statistics indicate that while the total rice harvest has grown at an average rate of 4.2% from 2007 to 2016, there are several abiotic and biotic stresses that restrict rice production, including rice blast disease (USDA 2017, Mgonja *et al.,* 2016, Mutiga *et al.,* 2017). Annual crop losses due to rice blast disease, caused by the fungus *Magnaporthe oryzae*, are reported to be up to 30%, with regional blast disease outbreaks causing up to 80% yield losses (Nalley *et al.,* 2016). Another major challenge in enhancing rice production is that smallholder farmers, who cannot afford the cost of fungicides, are the major growers of rice (Saito *et al.,* 2013; Nalley *et al.,* 2016). Therefore, farmer-friendly technologies such as breeding for rice blast resistance and surveillance of the fungal pathogen are urgently needed (Mutiga *et al.,* 2017).

To breed for blast resistant rice, an understanding of the pathology and molecular interactions between the fungus and its host is required. Knowledge of the pathosystem is necessary and involves information of different lifestyles that fungi can adopt and their co-evolution with plants, which also reflects their biological diversity and worldwide distribution (Occhipinti, 2013). Owing to adaptations of both the pathogen and crop plants, plant-pathogen interactions differ significantly across agro-ecological systems. Building on the many studies that have begun to unravel the genetic basis of plant-fungal interactions, current work is aimed at identifying the molecular signatures for virulence (Burdon and Thrall, 2009). The rice blast pathosystem is a long-standing co-evolutionary model system, however, the molecular interactions between *M. Oryzae* and rice has not been well studied using isolates from sub-Saharan Africa.

Hemibiotrophic fungal phytopathogens, such as *M. oryzae,* secrete effector proteins that counter plant defense signals and are key determinants of pathogenesis and virulence (Kamoun, 2007; Dodds *et al.,* 2009). Genomics-based effector discovery has identified genomic regions encoding effectors (Gibriel *et al.,* 2016). Plants in turn, exhibit recognition mechanisms that trigger production of resistance (*R*) proteins that directly, or indirectly, interact with effector molecules also called avirulence (*Avr*) proteins (Dodds and Rathjen, 2010; Petit-Houdenot and Fudal 2017). The activation of Effector-Triggered Immunity (ETI) in plants occurs upon the recognition of a pathogen *Avr* protein leading to a hypersensitive reaction (HR), which causes localized cell death, hence blocking disease progression (Jones and Dangl, 2006). This knowledge of *R*-*Avr* gene interaction is frequently utilized by scientists in breeding for superior crops that possess novel dominant resistance genes that cannot be infected by resident pathogen races. Genetic control of blast disease therefore involves development of resistant rice cultivars that harbor major *R* genes (Petit-Houdenot and Fudal, 2017). There is however a significant risk of resistance breakdown because of the selection for *Avr* genes to mutate leading to loss of recognition by *R* gene products. To overcome this risk, a new strategy is to pyramid multiple *R* genes into a locally adapted rice variety (Mutiga *et al.,* 2017; Pilet-Nayel *et al.,* 2017). More than 100 rice blast *R* genes have been identified in rice to date (Sharma *et al.,* 2012) and many of these have been cloned and characterized. The rice panel used in this GWAS includes lines harboring six major *R* genes (*Pia, Pita, Pi9, Pik, Pizt, Piz-5*) that have been functionally characterized and the *R* gene *Pi3* that is organized into a gene cluster with *Pii* (Wu *et al.,* 2015).

The advent of Next Generation Sequencing (NGS) and availability of fungal genomes has led to accelerated identification of *Avr* genes (Petit-Houdenot and Fudal 2017). However, due to exertion of selection pressure by *R* genes, fungal pathogens very frequently become virulent through evolution of *Avr* genes. In such cases, virulence can be achieved by inactivation, down-regulation or complete deletion of the *Avr* gene, or simply generating point mutations such as single nucleotide polymorphisms (SNPs) that disable recognition (Guttman *et al.,* 2014; Jones and Dangl, 2006). Accessibility of whole genomes or reduced representation of genomes has enabled genome-level analysis of plant pathogens. Sequence variation is often used to identify complex traits in plants, animals and microorganisms. There are several high-throughput methods that combine NGS with reduced representation of genomes (Glaubitz *et al.,* 2014). Genotyping-by-sequencing (GBS) is a simple, robust multiplex method that generates large numbers of SNPs at very low cost (Elshire *et al.,* 2011). GBS has been mostly used to study host plants (Torkamaneh *et al.,* 2017; Fernandez-Mazuecos *et al.,* 2017; Hussain *et al.,* 2017) because of the emphasis on resistance, however this technique can equally be applied to a pathogen. There are several combinatorial factors that can complicate the characterization of virulence in plant-pathogenic fungi, including genetic diversity present in natural populations, difficulty in conducting controlled crosses among genotypes, complexity of inheritance, inaccessibility to genomic information and the high costs associated with whole genome sequencing (Leboldus *et al.,* 2015). Using low cost GBS data can circumvent some of these issues (Leboldus *et al.,* 2015), and recent studies using GBS-based identification of quantitative trait loci in fungal pathogens have begun to emerge (Norelli *et al.,* 2017). GBS can be used to study population diversity and conduct Genome-wide Association Studies (GWAS) (Mjonga *et al.,* 2017).

GWAS is a potent tool used to detect genomic regions associated with natural variation in biological systems. This method is increasingly being used to study variation in plants, especially to fine map resistance genes in plants. However, GBS has only been sparingly used to detect genetic variants associated with pathogenicity/virulence in pathogens (Sanchez-Vallet *et al.,* 2017). In the last couple of years, GWAS has been utilized for successful identification of a wide range of alleles and candidate genes associated with disease or pathogenicity factors/phenotypes (Plissonneau *et al.,* 2017). There are more than 35 GWAS that have used SNPs as genetic markers for identifying genomic regions associated with plant response to pathogen infection (Bartoli and Roux, 2017) but only six of those GWAS reports on plant pathogens identified candidate pathogenicity determinants/genes. This includes one study of the bacterial pathogen *Pseudomonas syringae* (Monteil *et al.,* 2016) and five GWAS on fungal phytopathogens; *Heterobasidion annosum s.s*., *Parastagonospora nodorum, Fusarium graminearum, Puccina triticina and Zymoseptoria tritici* (Dalman *et al.,* 2013; Gao *et al.,* 2016; Talas *et al.,* 2016, Wu *et al.,* 2017, Hartmann *et al.,* 2017). These studies were conducted to directly identify loci in the genome of the pathogen contributing to virulence.

Our previous research (Mutiga *et al.,* 2017) on the virulence spectrum and genetic diversity of *M. oryzae* samples from SSA suggested that regional breeding strategies would be required for East and West Africa based on the observed associations between genetic relatedness and virulence spectrum. The objectives of the current study are as follows;

1. To dissect the pathogen-host interaction for African collections of *M. oryzae* based on inoculations on genotypes carrying known *R*-genes
2. To discover new avirulence genes, which could be useful in providing insights onto how *M. oryzae* interacts with rice
3. To gain insight into pathogen-host co-evolution; through knowledge of shared virulence genes; to assist the team in developing a breeding strategy to achieve durable resistance to rice blast disease

Here we use GBS and GWAS to detect SNPs to identify loci and putative genes as markers of virulence profile. To the best of our knowledge, this is the first study to report GWAS of *M. oryzae* isolates from sub-Saharan Africa. Natural isolates of *M. oryzae* tend to be female sterile (Ray *et al.,* 2016) and thus, this is also the first report of GWAS of an asexually reproducing fungal pathogen.

## MATERIALS AND METHODS

### Fungal Population used in the study

The genotyped *M. oryzae* population comprise 160 isolates from nine African countries (*n=128*) including Tanzania, Kenya and Uganda from East Africa, Nigeria, Burkina Faso, Benin, Togo, Mali and Ghana from West Africa, and international isolates (*n=32*) from seven major rice producing countries of China, India, Egypt, Philippines, South Korea, Colombia and United States (Table 1). The isolates were provided by D. Tharreau (Centre de Coopération Internationale en Recherche Agronomique pour le Développement, CIRAD, Montpellier, France), and by collaborators in the rice blast project which was supported through the Sustainable Crop Production Research for International Development (SCPRID) initiative funded by Bill & Melinda Gates Foundation through the Biotechnology and Biological Sciences Research Council (BBSRC) and Department for International Development (DFID) of the United Kingdom between 2012 and 2014.

**Table 1:**
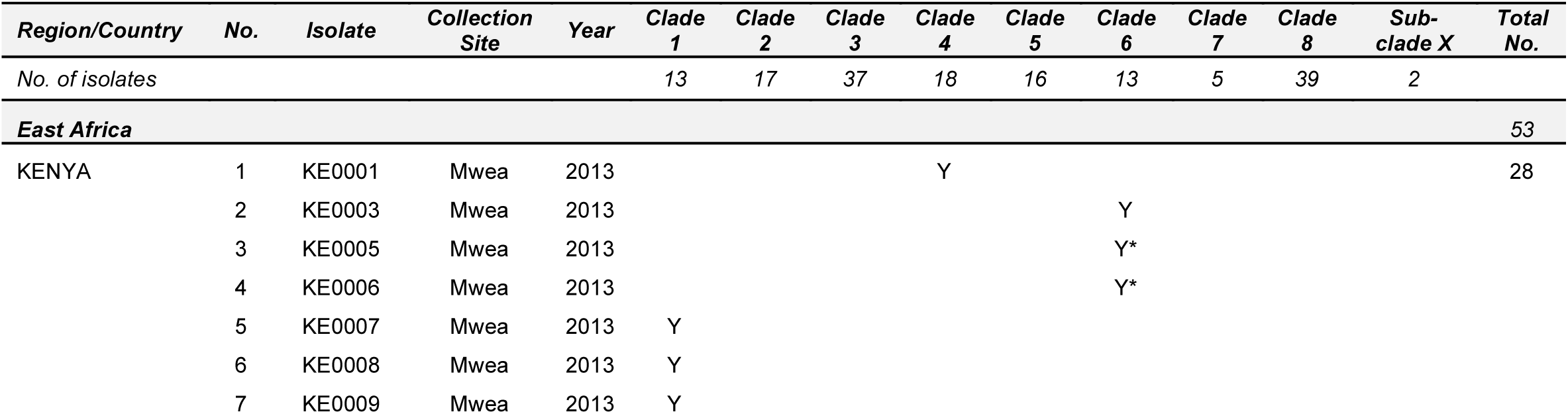
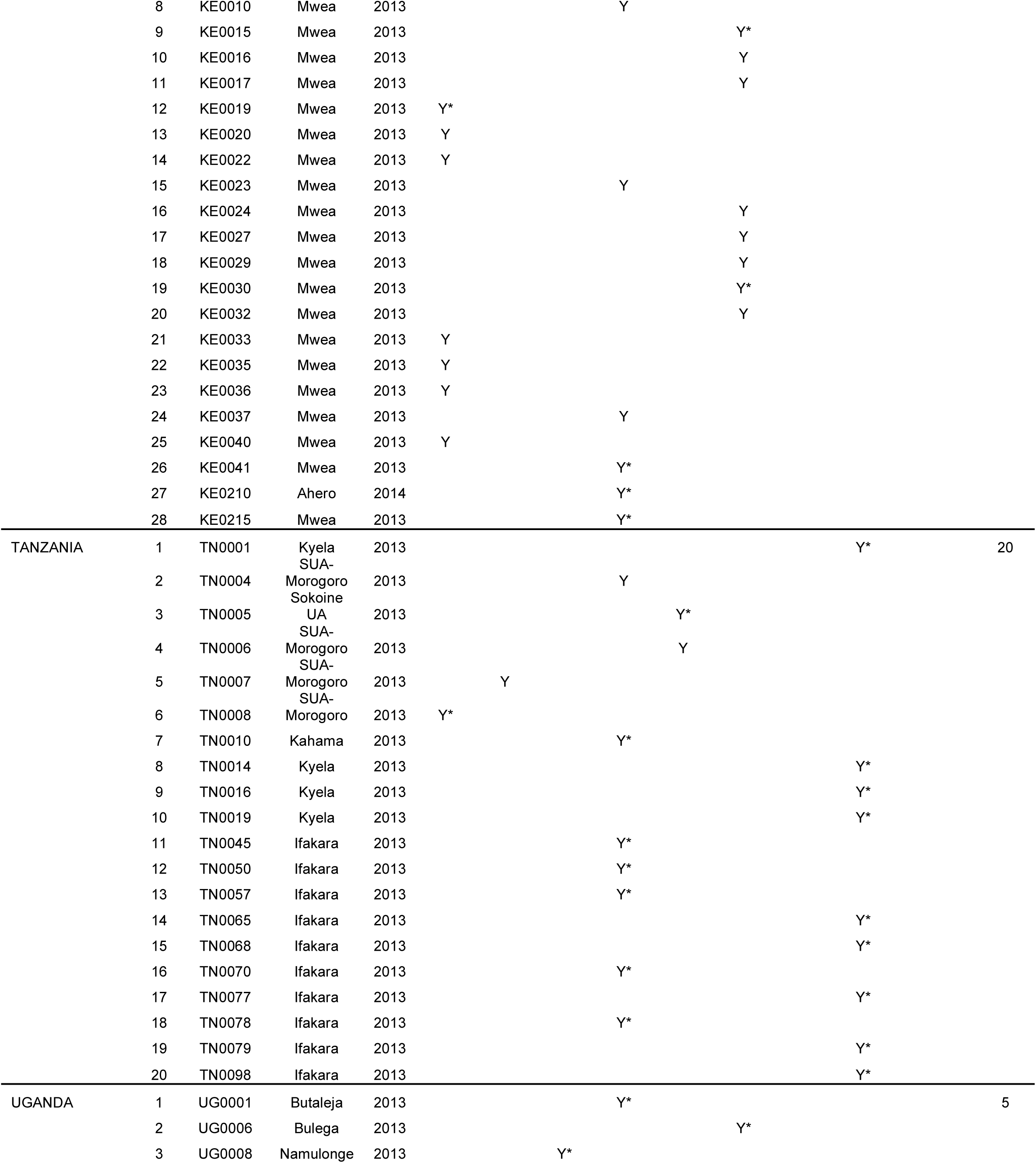
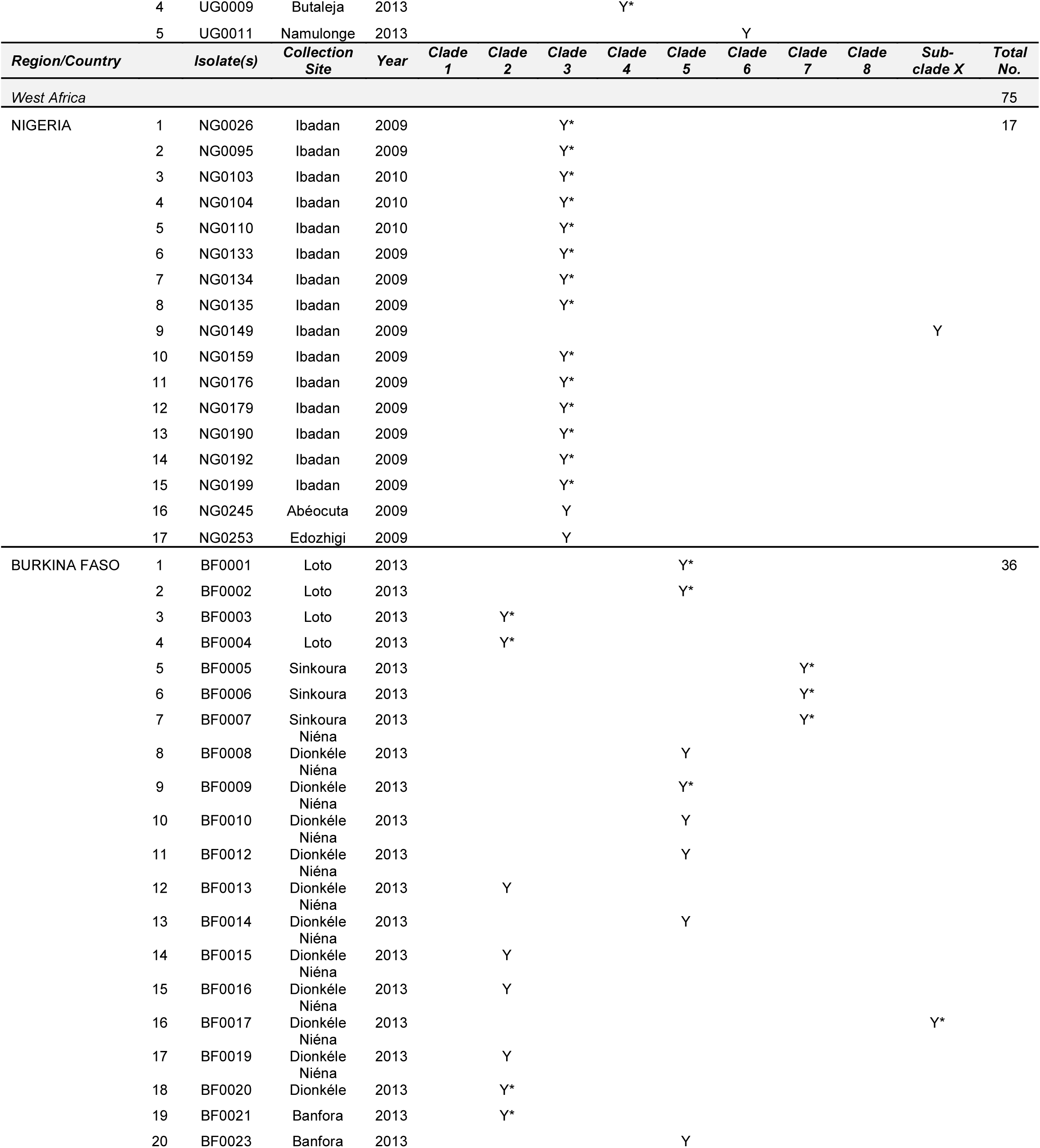
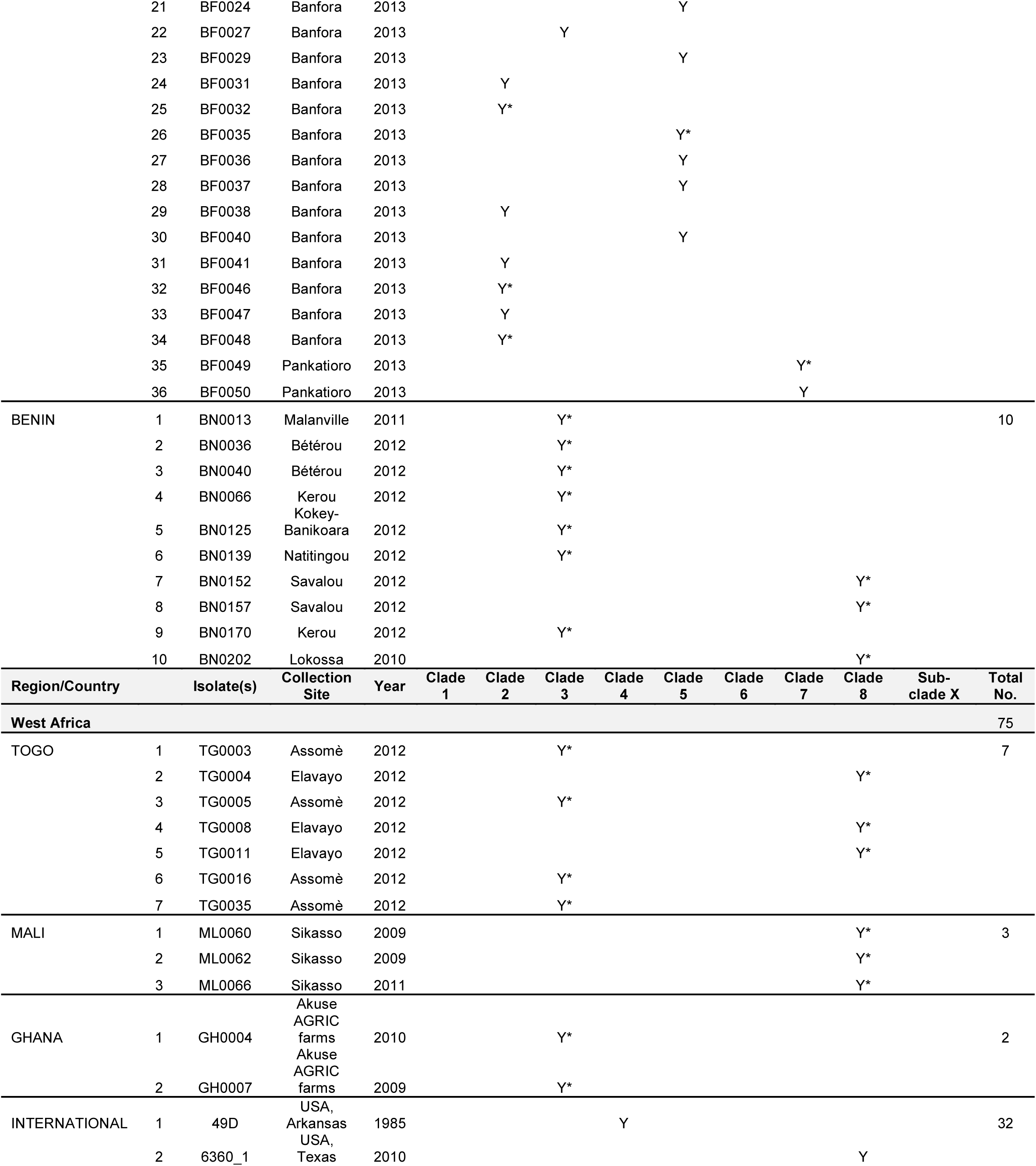
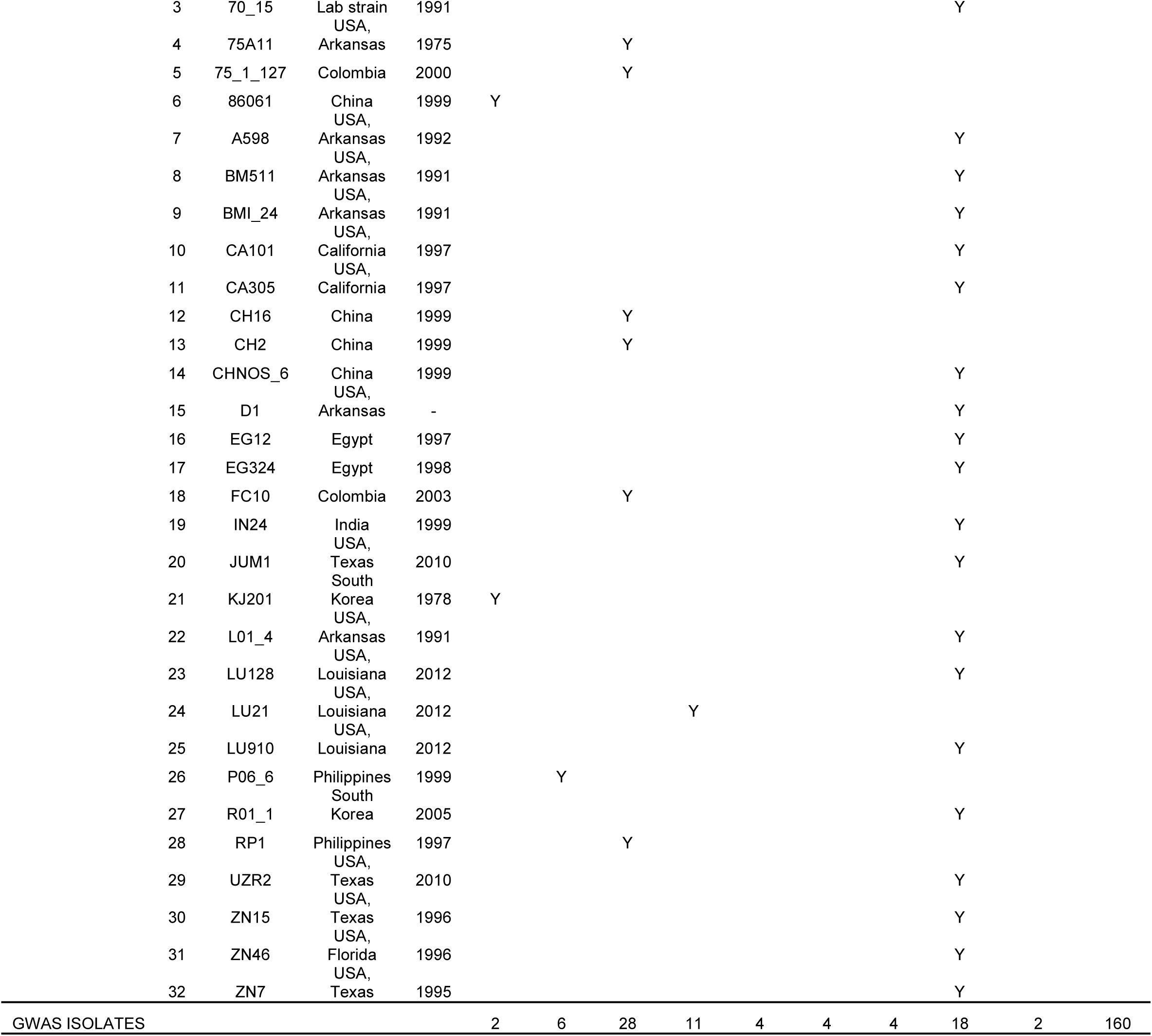
Country origins and collection sites of M. oryzae isolates from different parts of rice growing SSA countries and International regions (n=160). Y indicates the presence of the isolate in a clade and the asterisk represents isolates randomly chosen for GWAS (n=78).

### Fungal culture preparation, genomic DNA extraction, GBS library construction and Pathogenicity assays

Fungal isolates were grown in liquid PD broth and DNA was extracted using DNeasy Plant Mini Kit (Qiagen GmbH). GBS library construction, MiSeq sequencing and pathogenicity assays were conducted as previously described in Mutiga et al., 2017. A visual disease rating scale of 0-9 was used to assess foliar blast in pathogenicity assays.

### Rice Germplasm used in the study

Sixteen rice cultivars were used in the pathotyping assays. Differential rice lines IRBLA-a, IRBLTA CP1, IRBL3-CP4, IRBL9-W, IRBLK-KA, IRBLZT-T, IRBLZ5-CA(R), IRBLTA CT2 (n=8), African Upland Rice NERICA2 and NERICA5, African lowland rice FKR62N, African intraspecific lines TS2 and F6-36, susceptible check UZROZ275, *Pi9* donor line 75-1-127 and African *Oryza glaberrima* rice AR105. Nine of these cultivars had the presence of known blast resistance genes *Pia, Pita, Pi3, Pi9, Pizt, Piz-5, Pita* (Table 3). Rice cultivars containing the R-genes are known to be in the common background of an Asian *japonica* rice cultivar called Lijiangxituanheigu (LTH). The *Pi9* donor line 75-1-127 was kindly provided by the Wang lab (Plant Pathology, The Ohio State University), in collaboration with Liu *et al*., 2002. IRBLs (IRRI-bred blast-resistant lines) were provided by IRRI and bulked at DBNRRC, Stuttgart, Arkansas. UZROZ275 was provided by Correll lab (University of Arkansas). AR105, F6-36 and NERICA lines were provided by Dr. Ouédraogo (INERA-Burkina Faso, Linares 2002), NERICA lines were established by Africa Rice Center utilizing crosses of *O.glaberrima* and *O.sativa.*

**Table 2:** Description of isolates used in this study (a) Percentage of isolates from different SSA countries randomly chosen for GWA analyses (b) Percentage of isolates randomly chosen from each clade for GWA analyses.

| Region | Country | GWAS | Total isolates | % |
| --- | --- | --- | --- | --- |
| East Africa | Tanzania | 17 | 20 | 15 |
|  | Kenya | 7 | 28 | 35 |
|  | Uganda | 4 | 5 | 76 |
| West Africa | Nigeria | 14 | 17 | 61 |
|  | Burkina Faso | 14 | 36 | 25 |
|  | Benin | 10 | 10 | 31 |
|  | Togo | 7 | 7 | 80 |
|  | Mali | 3 | 3 | 46 |
|  | Ghana | 2 | 2 | 50 |
| Total |  | 78 | 128 |  |

(b)
| Clade | GWAS | Total isolates | % |
| --- | --- | --- | --- |
| 1 | 17 | 20 | 85 |
| 2 | 7 | 28 | 25 |
| 3 | 4 | 5 | 80 |
| 4 | 14 | 17 | 82 |
| 5 | 14 | 36 | 39 |
| 6 | 10 | 10 | 100 |
| 7 | 7 | 7 | 100 |
| 8 | 3 | 3 | 100 |
| X | 2 | 2 | 100 |
| Total | 78 | 128 |  |

**Table 3:** Description of rice cultivar panel used for GWAS and results of association analysis.

| Rice cultivar | Description | Target R-gene | Donor Line/pedigree | Association |
| --- | --- | --- | --- | --- |
| 1. IRBLA-a | Differential Line | *Pia* | Aichi Asahi | Yes |
| 2. IRBLTA CP1 | Differential Line | *Pita* | *C101PKT* | Yes |
| 3. IRBL3-CP4 | Differential Line | *Pi3* | *C104PKT* | Yes |
| 4. IRBL 9-W | Differential Line | *Pi9* | *WHD-1S-75-1-127* | Yes |
| 5. IRBLK-KA | Differential Line | *Pik* | *Kanto 51* | Yes |
| 6. NERICA5 | African upland | - | *WAB450-11-1-1-P31-HB* | Yes |
| 7. IRBLZT-T | Differential Line | *Pizt* | *Toride-1* | No |
| 8. IRBLZ5-CA (R) | Differential Line | *Piz-5* | *C101A51* | No |
| 9. IRBLTA CT2 | Differential Line | *Pita* | *C105TTP2L9* | No |
| 10. NERICA2 | African upland | - | *WAB450-11-1-P31-1-HB* | No |
| 11. FKR62N | African lowland | - | *WAS122-IDSA-1-WAS-6-1* | No |
| 12. TS2 | African intraspecific | - | NA | No |
| 13. F6-36 | African intraspecific | - | NA | No |
| 14. UZROZ275 | susceptible check | - | NA | No |
| 15. 75-1-127 | *Pi9* donor line | *Pi9* | NA | No |
| 16. AR105 | *Oryza glaberrima* (African) | - | NA | No |

### Phylogenetic and PC analysis

The high-quality filtered SNP panel consisting of 617,281 SNPs was used to conduct the neighbor-joining phylogenetic cluster and principal component analyses using TASSEL v5.2.1.6 (Bradbury et al., 2007). The unrooted neighbor-jointing tree was constructed using the calculations derived from pairwise distance matrix between individuals/taxa. PC analysis was performed and the values of first three components were exported to R for plotting the 3D plot using P3D function (R core team 2015)

### Genome-wide association analysis

Association between SNPs (GBS) and disease scores (pathogenicity assay) data was implemented using TASSEL v5.2.3 (Bradbury et al., 2007). GWA analyses were named after the cultivar of rice from which the disease scores were obtained. A Generalized Linear Model (GLM) was used to execute the association analysis utilizing the least squares fixed effects linear model. The *p-*values obtained from the analysis were visualized in a Q-Q plot using the R package “qqplot” (R core team., 2013). *p*-value cutoffs differed each analysis as they were dependent on the deviation of *p*-values from X=Y line representing true associations.

### Identification of putative virulence-related genes

A region 500bp upstream and downstream surrounding the SNPs was extracted from the respective contigs using a customized R script (R Core team., 2017). The size of the sequence determined was based on the average length of a protein sequence in NCBI db. The homology of the sequence surrounding the SNPs was identified using the DNA database from NCBI (v 2.7.1, 23 Oct, 2017).

## RESULTS

### Geographical representation of fungal isolates from sub-Saharan Africa and the major rice growing areas of the world

The field isolates used for both pathotyping and genotyping in this study (*n=160*) were collected from blast-infected rice plants from nine countries of sub-Saharan Africa, including East African countries (Kenya, Tanzania and Uganda), and West African countries (Benin, Burkina Faso, Ghana, Mali, Nigeria, and Togo, as shown in Fig. 1A. In addition, some isolates from major rice growing areas outside of SSA, such as China, India, Egypt, Philippines, Colombia, South Korea and United States, were included as comparators in the analysis. SSA isolates represent the major regions as follows: those from West African countries were the highest (*n=75*, 46.9%) compared to East African (*n=53*, 33.1%) and International isolates (*n=32*, 20%) (Table 1). There was variable sampling within countries and Burkina Faso had the largest sampling size (*n=36*, 22.5%) while the least sampling size was from Ghana (*n=2*, 1.25%).

**Figure 1:**
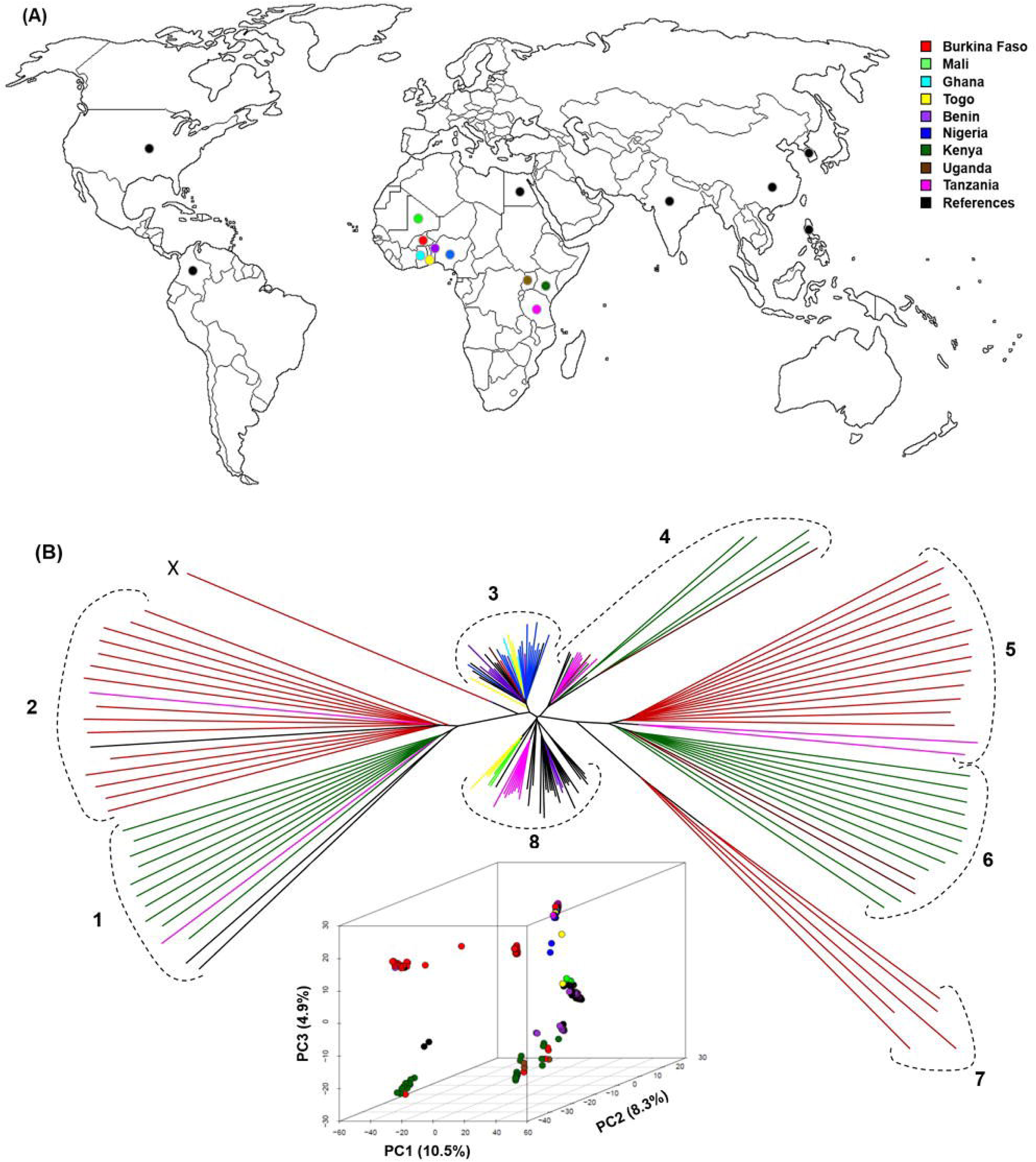
Genetic Diversity of *M. oryzae* isolates from sub-Saharan Africa (SSA) (A) Geographic origin of SSA field solates and Internat onal comparator isolates. (B) Unrooted neighbor joining tree of *M. oryzae* isolates *(n=160)* based on pair wise distance matr x calculated by TASSEL v5.0 (C) 3D Pr ncipal Component Analysis scatter plot show ng the first three components (PC1, PC2, and PC3) Colored lines and colorfilled circles represent SSA countr es, black lines and filled black circles represent International isolates (Table 1).

### GBS identifies genetic diversity among SSA field isolates

Genotyping of 160 isolates of *M. oryzae* from diverse geographic regions generated a 617K (617,281) panel of polymorphic markers. SNPs were then used to assess diversity of isolates using unrooted neighbor-joining trees based on a pair-wise distance matrix. The analyses showed segregation of isolates into 8 clades and 1 independent clade X. Each cluster of isolates belonged to the same country or region (Fig. 1B and C). Most of the East African countries grouped into clades 1, 4, 6, and 8 and West African countries grouped into clades 2, 3, 5, 7, and 8. Two isolates, one from Burkina Faso (BF0017) and another from Nigeria (NG0149), segregated from the rest of the major clades as an independent subclade X.

Isolates from the West African country of Burkina Faso clustered into 3 clades (called 2, 5 and 7) although some isolates from the East African country Tanzania were also present in these clades. Kenyan isolates segregated into 2 predominant clades (1 and 6), isolates in Clade 1 clustered with 2 international isolates (from the USA and South Korea) along with a Tanzanian isolate. Seven independent Kenyan isolates clustered in Clade 4 with Tanzanian, Ugandan and 2 isolates from the United States. Clade 3 members were mostly from the West African country of Nigeria, which clustered closely with isolates from other West African countries of Benin, Togo, Ghana and Burkina Faso, an isolate from East African country of Uganda and the international isolates from USA, China, Colombia and Philippines. Clade 8 had the largest number of isolates at 39, and its subclades mostly showed clustering of isolates within each country. Isolates from Tanzania were, for example, more similar to the international isolates 70-15, CHNOS-06 and BMI-24, whereas 3 isolates from Benin sub-clustered with international isolates from the USA and Egypt. Isolates from Mali and Togo were similar to the single Indian isolate IN24.

To explore the similarity among isolates in greater detail, we conducted Principal Component Analysis (PCA) and constructed a 3D scatter plot based on the discriminatory power of individual SNPs and used this to identify population structure patterns within West African and East African isolates, (Fig. 1C). The first three principal components (PCs) explained 23.7% of the total variance observed in the dataset. The scatter plot based on the first 3 PCs also corresponded to a close phylogenetic relationship between the isolates wherein, they segregated into 8 groups mostly based on country or region. While PC1 accounted to 10.5% of the variability, PC2 and PC3 accounted for 8.3% and 4.9% of the variability respectively. The clustering of isolates observed in the unrooted neighbor joining tree was consistent with the PC analysis.

### Description of isolates chosen for GWAS analysis

Representative isolates from different SSA countries were selected based on the distinct clades they occupied based on phylogenetic analysis and the availability of pathotyping data for GWAS analysis. A total of 78 isolates were used for the analysis, which accounts to 49% of the complete dataset and 80% of SSA dataset. All of the isolates from the clades 6, 7, 8 and “X” were used for GWAS, whereas for clades 1,2,3,4 and 5, 85%, 25%, 80%, 82.3% and 38.8% of the isolates were analyzed as shown in Table 2. International isolates were not included in the GWA analysis.

### GWAS identifies genomic markers of virulence

Principal Component Analysis (PCA) has been used as an alternative to population structure analysis for studying population stratification from genotypic data (Patterson et al., 2006). In this study, we used PCA scores from the genotypes as covariates in the GWAS analyses. GWA analyses were conducted to identify genomic locations/putative genetic loci associated with the ability to cause rice blast disease in a given host cultivar. A Generalized Linear Model (GLM) was used to perform GWA using least squares fixed effects linear analysis using SNPs derived from GBS and maximum score from the disease rating observed. From the 16 GWA analyses conducted, six showed association with disease and 10 of them showed no association (Table 3). Most *p*- values observed in the scatter plot with association to disease were similar to the expected diagonal in the QQ-plot, demonstrating the appropriateness of the GLM Model for GWAS used in this analysis (see Fig. 2).

**Figure 2:**
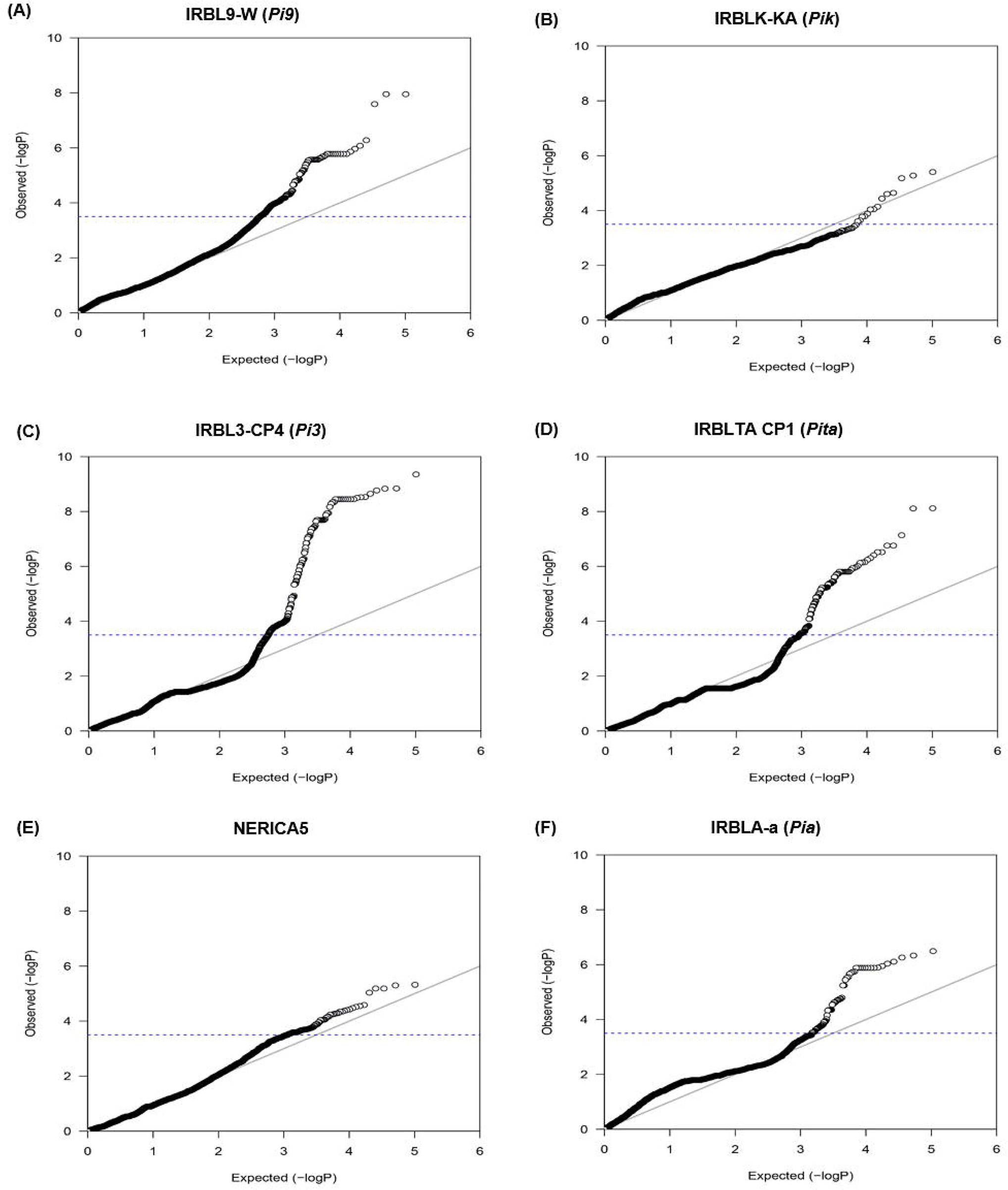
Quantile-Quantile plots for the 6 GWA analyses using a generalized linear model (GLM). The x-axis corresponds to expected values of negative logarithm of *P* and y-axis corresponds to observed values of negative logarithm of *P.* (A) Rice blast differential rice line IRBL9-W harboring the R-gene *Pi9* (B) Rice blast differential rice line IRBLK-KA harboring the R-gene *Pik* (C) Rice blast differential rice line IRBL3- CP4 harboring the R-gene *Pi3* (D) Rice blast differential rice line IRBLTA-CP1 harboring the R-gene *Pita* (E) African upland rice cultivar NERICA5 **(F)** Rice blast differential rice line IRBLAa harboring the R-gene *Pia.* Blue dotted lines indicate –*logP* cutoff

Based on the Q-Q plot analysis of each GWA analysis, independent observed – log*P* cutoffs were then used to obtain SNPs associated with the ability to cause disease. A cutoff of 3.5 was used in the analyses of IRBLTA, IRBL3 and NERICA5, which yielded 95, 173 and 64 associated markers, respectively. A cutoff of 4.5 was used for the analyses of IRBLA-a and IRBL9-W, which yielded 135 and 54 markers, respectively, while a cutoff of 4.0 was used for IRBLK-KA which generated 7 associated markers (Fig 3). A total of 528 markers were obtained from the 6 GWA analyses. However, due to the absence of a completely annotated Guy11 genome, a *de novo* assembly of Guy11 contigs was utilized (Source: Darren Soanes, University of Exeter) as the reference genome in the alignment process during GBS analysis. This did not provide chromosome locations and SNP positions on the genome, but did provide SNP locations in the *de novo* contigs. We then used the contig information, coupled with the SNP marker location, to identify the genomic location of markers compared to the published 70-15 *M. oryzae* genome (Dean et al., 2005).

**Figure 3:**
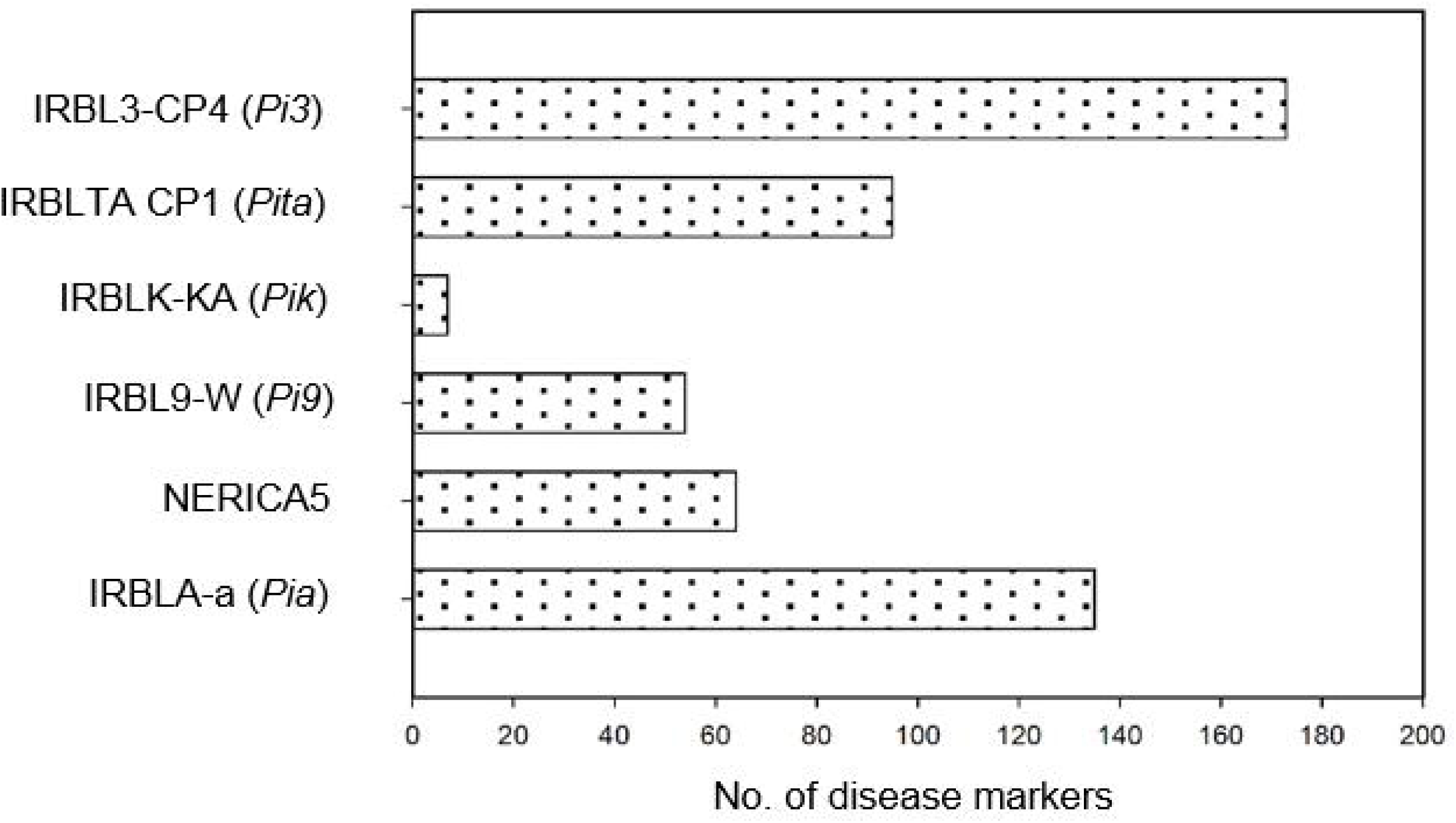
Number of rice blast disease-associated SNP markers identified in individual GWA analyses. Fungal isolates show virulence on the respective rice cultivars shown. Five cultivars carry single major blast disease resistance genes

### GWAS guides prediction of virulence-associated genetic loci

A nucleotide sequence similarity search using NCBI BLASTn database (Altschul et al., 1990) was conducted with the sequences encompassing the identified SNPs, (500bp upstream and downstream based on average length of protein sequence in NCBI db; 325 amino acids long) associated by GWAS with the ability to cause disease on a given host. Their homologies were categorized into four different types, which are SNP markers that are located in (a) predicted genes (PG), (b) hypothetical proteins (HP), (c) Other (OT); those present in non-coding genomic sequences, such as repeat regions, (d) no similarity (NS) (Table S1-S6).

The highest number of SNPs associated with blast disease were observed in the GWA analysis involving rice line IRBL3-CP4. This yielded 173 markers, out of which a majority of the markers (*n=71*, 41%) were predicted to be hypothetical proteins, uncharacterized ORFs whose structural genes exist but where corresponding translation products are so far unidentified. Fifty two of the markers (30%) were identified as predicted genes with known functions, and 11 genes (6.35%) reported in other uncharacterized genomic regions or repeat regions. The sequence location of 18 markers (10.4%) showed no similarity to any sequence present in the NCBI Nucleotide database (Fig 4).

**Figure 4:**
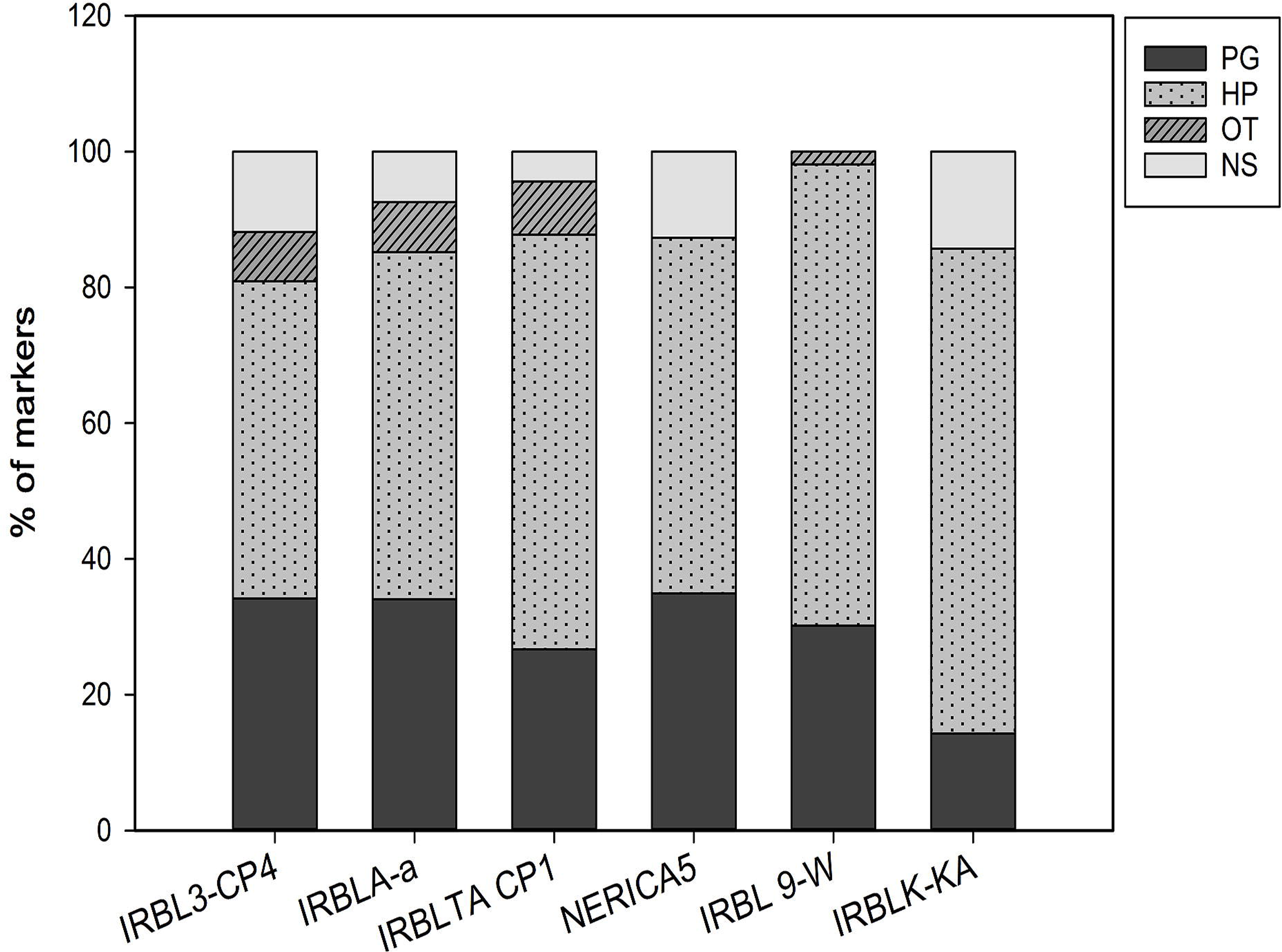
Distribution of SNP marker locations in different GWA analyses as predicted by NCBI’s DNA database. PG predicted genes; HP hypothetical protein; OT other, present in genomic sequences, repeat regions on genome; NS no similarity.

The NERICA5 GWA analysis had the highest percentage of predicted gene similarity (34.9%), while the IRBLK-KA had the lowest at 14.2%. Most of the SNP markers were located in hypothetical protein-encoding genes across all the studies, with an average of 58.4% (+/-4.05) showing this association. Some markers from IRBL3-CP4, IRBLA-a, IRBLTA-CP1 and IRBL-W analyses had similarities with fosmid clone genomic regions or repeat regions and this was predominant in 3 analyses (IRBL3-CP4, IRBLA-a, IRBLTA-CP1) accounting to only 7% of the markers. In some cases, the DNA sequence in which some of the markers were present were dissimilar to sequences in the *nt* database and this accounted to 11.8% in IRBL3-CP4, 7.4% in IRBLA-a, 4.4% in IRBLTA-CP1, 12.6% in NERICA5 and 14.2% in IRBLK-KA (Fig. 4). Overall, sequence similarity to hypothetical protein-encoding genes was higher compared to predicted genes.

### Consistent SNPs associated with virulence of M. oryzae

A closer look at predicted genes across different GWA analyses indicated the recurrence of specific genes. For example, a AGC/RSK protein kinase (*MGG_07012*), a class of protein previously implicated in regulating signaling events that coordinate growth and morphogenesis in fungi (Lee *et al.,* 2016) and a rRNA processing protein FCF2 (*MGG_00473*) involved in ribosome biogenesis (Rempola *et al.,* 2006), were reported in five of six GWA analyses conducted (Table 4).

**Table 4:**
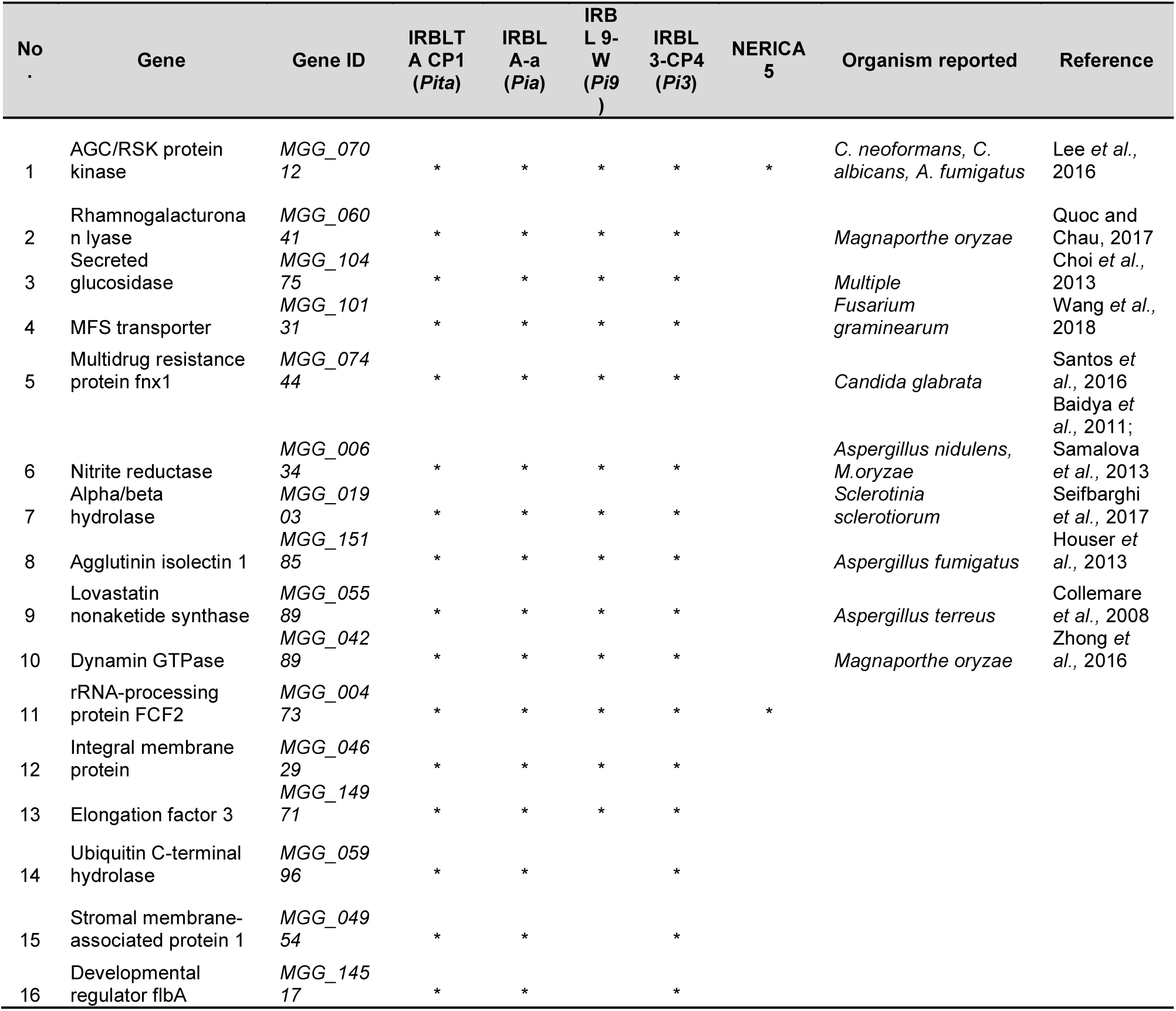
Putative virulence genes (*n=16*) that occurred in the five rice blast disease associated GWA analyses.

IRBLTA CP1, IRBLA-a, IRBL9-W and IRBL3 CP4 showed recurrence of 11 genes. (1) Rhamnogalacturonan lyase (*MGG_06041*) (2) Secreted glucosidase (*MGG_10475*) (3) MFS (Major facilitator Superfamily) transporter (*MGG_10131*), (4) Integral membrane protein (*MGG_04629*) (5) Multidrug resistance protein fnx1 (*MGG_07444*) (6) Nitrite reductase (*MGG_00634*) (7) Alpha/beta hydrolase (*MGG_019030*) (8) Agglutinin Isolectin 1 (*MGG_15185*) (9) Elongation Factor 3 (*MGG_14971*) (10) Lovastatin nonaketide synthase (*MGG_05589*) (11) Dynamin GTPase (*MGG_04289*) (Table 3). Three genes were identified in IRBLTA CP1 (*Pita*), IRBLA-a (*Pia*) and IRBL3-CP4 (*Pi3*), they are (1) ubiquitin C-terminal hydrolase (*MGG_05996*) (2) Stromal membrane-associated protein 1 (*MGG_04954*) and (3) the Developmental regulator flbA (*MGG_14517*). DNA-repair protein rad13 (*MGG_00155*) was reported in the three analyses of IRBLTA CP1 (*Pita*), IRBLA-a (*Pia*) and IRBL-9W (*Pi9*) (Table 4).

### Multiple SNPs in Genes/Genomic regions associated with virulence

Multiple SNPs in a region/gene were observed across 5 analyses (IRBLTA CP1, IRBL3 CP4, IRBLA-a, IRBLA 9-W and NERICA5). These SNPs were found to be located in hypothetical protein-encoding genes, genes with predicted products and other regions in the genome (Table 5). IRBLTA CP1 had the highest number of genes (*n=12*), showing multiple SNPs, while NERICA5 had the lowest (*n=1*). The IRBL3 CP4 had the second highest number (*n=9*) and the analyses of IRBLA-a and IRBL9-W had similar number of genes with multiple SNPs (*n=3*). SNPs varied in number from a maximum of nine to a minimum of two SNPs per region/gene. Seven genes/regions harboring multiple SNPs were commonly present in one or more of the GWA analyses. *MGG_16548* was identified in IRBLTA CP1, IRBL3 CP4, IRBL 9-W and NERICA5 while *MGG_07814* and *MGG_10232* were identified in IRBLTA CP1, IRBL3 CP4 and IRBL 9-W. Y34 repeat region of *M. oryzae* with accession JQ929669 was identified in IRBLTA CP1 and IRBL3 CP4, the fosmid SK2054 genomic sequence was found in IRBLTA CP1 and IRBLA-a, *MGG_02369* was identified in IRBL3 CP4 and IRBLA-a and *MGG_17298* was present in three of the GWA analyses of IRBLTA CP1, IRBL3 CP4 and IRBLA-a.

**Table 5:** Genes/genomic regions with multiple SNPs associated with rice blast disease. Highlighted genes were identified in five of the GWA analyses.

| Study | No. | Gene/Locus | Homology | No. of SNPs |
| --- | --- | --- | --- | --- |
| IRBLTA CP1 (Pita) | 1 | **MGG_10232** | *Magnaporthe oryzae* 70-15 hypothetical protein mRNA | 9 |
|  | 2 | **MGG_07812** | *Magnaporthe oryzae* 70-15 hypothetical protein partial mRNA | 4 |
|  | 3 | **JQ929669** | *Magnaporthe oryzae* strain Y34 repeat region | 4 |
|  | 4 | **MGG_16548** | *Magnaporthe oryzae* 70-15 hypothetical protein partial mRNA | 3 |
|  | 5 | **MGG_17298** | *Magnaporthe oryzae* 70-15 hypothetical protein partial mRNA | 4 |
|  | 6 | **KT599263** | *Magnaporthe oryzae* strain BM1-24 clone fosmid SK2054 genomic sequence | 2 |
|  | 7 | **MGG_01018** | *Magnaporthe oryzae* 70-15 hypothetical protein partial mRNA | 2 |
|  | 8 | **MGG_02059** | *Magnaporthe oryzae* 70-15 hypothetical protein mRNA | 2 |
|  | 9 | **MGG_07812** | *Magnaporthe oryzae* 70-15 hypothetical protein partial mRNA | 2 |
|  | 10 | **MGG_04289** | *Magnaporthe oryzae* 70-15 dynamin GTPase partial mRNA | 2 |
|  | 11 | **MGG_14358** | *Magnaporthe oryzae* 70-15 hypothetical protein partial mRNA | 2 |
|  | 12 | **MGG_15795** | *Magnaporthe oryzae* 70-15 hypothetical protein partial mRNA | 2 |
| IRBL3 CP4 (Pi3) | 1 | **MGG_10232** | *Magnaporthe oryzae* 70-15 hypothetical protein | 8 |
|  | 2 | **MGG_07812** | *Magnaporthe oryzae* 70-15 hypothetical protein partial mRNA | 4 |
|  | 3 | **JQ929669.1** | *Magnaporthe oryzae* strain Y34 repeat region | 4 |
|  | 4 | **MGG_16548** | *Magnaporthe oryzae* 70-15 hypothetical protein | 3 |
|  | 5 | **MGG_17298** | *Magnaporthe oryzae* 70-15 hypothetical protein | 4 |
|  | 6 | **MGG_02369** | *Magnaporthe oryzae* 70-15 hypothetical protein | 4 |
|  | 7 | **MGG_00473** | *Magnaporthe oryzae* 70-15 rRNA-processing protein | 3 |
|  | 8 | **MGG_03564** | *Magnaporthe oryzae* 70-15 mitochondrial RNA-splicing | 4 |
|  | 9 | **MGG_17542** | *Magnaporthe oryzae* 70-15 hypothetical protein mRNA | 4 |
| IRBLA-a (*Pia*) | 1 | **MGG_17298** | *Magnaporthe oryzae* 70-15 hypothetical protein partial mRNA | 4 |
|  | 2 | **KT599263.1** | *Magnaporthe oryzae* strain BM1-24 clone fosmid SK2054 genomic sequence | 3 |
|  | 3 | **MGG_02369** | *Magnaporthe oryzae* 70-15 hypothetical protein partial mRNA | 4 |
| IRBL9-W (*Pi9*) | 1 | **MGG_10232** | *Magnaporthe oryzae* 70-15 hypothetical protein mRNA | 9 |
|  | 2 | **MGG_07812** | *Magnaporthe oryzae* 70-15 hypothetical protein partial mRNA | 3 |
|  | 3 | **MGG_16548** | *Magnaporthe oryzae* 70-15 hypothetical protein partial mRNA | 3 |
| NERICA5 | 1 | **MGG_16548** | *Magnaporthe oryzae* 70-15 hypothetical protein partial mRNA | 3 |

## DISCUSSION

This study utilizes SNPs derived from high-throughput genotyping using a GBS approach to characterize isolates of *M. oryzae* from Africa based on genetic diversity, and to then identify genomic regions associated with virulence on rice. While in-depth phylogenetic analysis of a sub-set of the isolates used in this study has been reported previously (Mutiga *et al.,* 2017), we report here the results of GWAS to define putative genes associated with virulence in isolates from SSA. The study involves rice lines with and without known blast resistance genes. The use of monogenic lines with known blast resistance genes was aimed at validating the known effector/avirulence genes, while the inclusion of the other lines, is expected to provide insight into the potential resistance genes and/or more insights in the pathosystem. This is the first study to identify genomic regions associated with virulence of *M. oryzae* isolates from sub-Saharan Africa based on GWAS. It is expected that these findings will boost knowledge of molecular Magnaporthe-rice pathosystem in sub-Saharan Africa, and hence enhance the breeding for durable blast resistance.

In this study, *M. oryzae* isolates were sampled from nine SSA countries, which were categorized into East and West Africa (Table 1). Rice cultivars commonly grown in this region were utilized in the pathogenicity assays to study virulence (Table 3). Efforts were focused on a better understanding of the Magnaporthe-rice pathosystem by studying population structure of the pathogen and conducting GWAS to identify novel markers associated with pathogenicity. Previously, a robust characterization of *M. oryzae* isolates from SSA was conducted using a combination of phylogeny of GBS-derived SNPs and pathogenicity assays using rice differential lines harboring known blast resistance genes (Mutiga *et al.,* 2017). GBS data phylogenetics revealed genetic relatedness of the pathogen collection from different West and East African regions. A comprehensive analysis of the genetic relatedness of 78 *M. oryzae* isolates from SSA showed a clear segregation of East and West African isolates compared to isolates from outside Africa (Mutiga *et al.,* 2017). The relationship between SNPs and virulence of the isolates was studied based on disease scores using standard least square regression and an evidence of association between genetic diversity and virulence of the isolates was identified. Analysis of the association between genetic relatedness and virulence showed that 77% of isolates in the three clades with highest mean disease score were from West African region known to have had a longer history of rice cultivation compared to East Africa. The emergence of avirulent and highly virulent strains may have resulted from continued rice production, breeding diverse genotypes and trade, resulting in a continuing modification in virulence spectrum of the isolates. It was suggested that the observed differences in virulence amongst the isolates clustered within independent clades conclude that some of these SNPs can lead us to the identification of genomic regions associated with virulence or avirulence. However, since that study used relatively few isolates, in this study the sampling size was increased to obtain a more holistic view of the population structure, augment the statistical power, as well as obtain meaningful associations by conducting GWAS analysis to identify novel virulence factors.

A GBS-SNP phylogeny of 160 isolates including 32 international isolates was conducted highlighting the clustering of isolates into eight unique clades (Fig 1). Segregation of West African and East African isolates into specific clades was clearly observed, which indicated the prevalence of higher variability and disease occurrence. Interestingly, no isolates from a single country were monophyletic, suggesting independent events of separate pathogen introductions in each country. This result is supported by the fact that *M. oryzae* is a globally dispersed trade pathogen (Tharreau *et al.,* 2009), and there is no clear country-specific geographic pattern of samples due to international trade and germplasm movement. Three isolates from Tanzania were, for example, closely related to most of the Burkina Faso isolates, suggesting that either of them could be the main source of disease outbreak within the region. Similarly, Burkina Faso isolates belonging to clades 2 and 5 are closely related to Kenyan isolates from clade 1 and 6. Although clade 8 was the largest-clade containing the majority of isolates (*n=39*), 53.8% of those are international strains, from the U.S.A., China, Egypt and India. Moreover, 23% were from the East African country of Tanzania that were related to isolates from West African countries of Togo (7.7%), Mali (7.7%) and Ghana (7.7%). The presence of admixture of isolates such as this, is indicative of global rice trade. With the exception of one isolate from the East African country of Uganda, clade 3 predominantly consists of isolates from the West African countries of Nigeria, Burkina Faso, Benin, Togo, Mali, Ghana with its origins from China, Philippines, Colombia and the U.S.A. Principal Component analysis was largely consistent with the phylogenetic analysis (Fig 1B and C).

To our knowledge, there have been only two other reports apart from this (Mutiga *et al.,* 2017) of a fungal study using GBS for population structure and genetic diversity analysis (Milgroom *et al.,* 2014, Rafiei *et al.,* 2018). Both of those studies focused on the ascomycete fungus *Verticillium dahliae* which reproduces mitotically and the population structure of which is highly clonal. GBS was performed on 141 *V. dahliae* isolates collected from diverse geographical and host origins that yielded 26,748 SNPs. The authors identified a large number of candidate SNPs distinct to lineages that can be used in the development of diagnostic markers, providing a strong suggestion that GBS can be used as a potential genotyping method for the analysis of clonally propagating fungi (Milgroom *et al.,* 2014). The current study yielded 617,281 filtered, high-quality SNPs that constitute a substantial SNP dataset in which the SNPs are evenly distributed throughout the genome, providing a more complete assessment of population structure, which to our knowledge is the first report of such a magnitude of SNPs in such a field study. The large panel of SNP markers used in this study will provide precise discrimination among geographical regions and enhance our association analysis of the Magnaporthe-Rice pathosystem.

The rice cultivar panel used in this study includes a diverse panel of rice lines (Table 3) including IRRI-bred blast-resistant lines (IRBLs) with known single target *R* genes, African rice germplasm (upland, lowland and intraspecific lines), a susceptible check and *Oryza glaberrima*, the African rice species that is one parent in the interspecific NERICA lines. This diverse panel was selected ensuring adequate sampling of isolates across different ecosystems that also covers screening of germplasm with known *R* genes.

A Generalized Linear Model was used in GWA analysis that utilizes a fixed effects linear model for association between segregating sites and phenotypes. It accounted for population structure using covariates, which indicates a degree of membership in underlying populations. Principal component (PC) scores obtained from PC analysis of genotypes were used as covariates, because there is no kinship in *M. oryzae* populations. Of the 16 GWA analyses, six showed association with disease in SSA, including differential rice lines that harbored the dominant *R* genes; *Pia, Pita, Pi3, Pi9* and *Pik* and the African interspecific upland rice variety NERICA5. Varying cutoffs for observed p-values were utilized to maximize number of markers associated with disease in each study (Fig 2). We obtained a significant number of markers (*n=528*) from the six studies (Fig 3), whose location on the genome was identified and NCBI BLASTn analysis confirmed the homology of sequences (500bp upstream and downstream) surrounding each SNP. The majority of those markers (n=269) were located in genes encoding hypothetical proteins (Fig 4) with uncharacterized ORFs the structural genes of which can be predicted, but where their analogous translation products remain unidentified. It has been reported that *M. oryzae* has more than 12,000 protein-encoding genes and 65% of them are not yet annotated (Li *et al.,* 2018). Thus, it is not surprising that a majority of the identified markers returned hypothetical protein hits. The second highest number of markers (*n=161*) were located in genes with predicted products and it can be noted that there are several genes that were repeatedly identified across 5 of 6 GWA studies (*n=16*) (Table 4). These are *MGG_07012, MGG_06041, MGG_10475, MGG_10131, MGG_07444, MGG_00634, MGG_01903, MGG_15185, MGG_05589, MGG_04289, MGG_00473, MGG_04629, MGG_14971, MGG_05996, MGG_04954* and *MGG_14517*. Some of these genes have reported to be associated with disease in Magnaporthe (*n=3*), other fungal systems (*n=7*) and unreported (*n=6*).

Out of the 16 genes that are considered as encoding putative virulence factors in this study, one of them, *MGG_07012*, a AGC/RSK kinase was consistently found across 5 of the 6 GWA analyses, including the African rice cultivar NERICA5, which suggests that this gene plays an important role in virulence of *M. oryzae* isolates from SSA. AGC kinases have been reported to be a common pathogenic protein kinase in fungal pathogens of humans, such as *Cryptococcus neoformans, Candida albicans* and *Aspergillus fumigatus* (Lee *et al.,* 2016). The AGC kinase subfamily contains 60 members including RSK. In humans these kinases have been shown to mediate important cellular functions and their mutation and/or dysregulation can cause human diseases (Pearce *et al.,* 2010). Furthermore, 9 genes were repeatedly found in four analyses [IRBLTA CP1 (*Pita*), IRBLA-a (*Pia*), IRBL 9-W (*Pi9*), IRBL3-CP4 (*Pi3*)], all of which have been previously reported to be involved in virulence of *M. oryzae* or other fungal systems. The 9 respective genes identified in SSA *M. oryzae* isolates are: (1) *MGG_06041*, a Rhamnogalactouronan lyase (Quoc and Chau, 2017) that recognizes and cleaves the ∞-1,4 glycosidic bonds in the backbone of rhamnogalactouronan-I, a major component of the plant cell wall polysaccharide, pectin. (2) the MFS transporter *MGG_10131*, transporters such as these have recently been shown to be important for production of mycotoxins such as deoxynivalenol (DON) in *Fusarium graminearum* (Wang *et al.,* 2018). DON plays a key role in infection of host plants (Wang *et al.,* 2018). (3) the Nitrite reductase (NR)-encoding gene *MGG_00634*. NR catalyzes the formation of Nitric oxide (NO) which can have implications in virulence of fungal pathogens by regulation of mycotoxin biosynthesis, as reported in *Aspergillus nidulans* (Baidya *et al.,* 2011). The requirement of NO for successful colonization of host by *M. oryzae* has been reported suggesting a critical role in appressorium formation (Samalova *et al.,* 2013). (4) Alpha beta-hydrolases similar to *MGG_01903*, identified in this study, were shown to be induced during infection in *S. sclerotiorum*, where some of them are involved in lipid degradation, as esterases, or lipases, and others act as hydrolytic enzymes in general (Seifbarghi *et al.,* 2017). (5) Secreted glucosidase *MGG_10475* was identified in four of the six GWA analyses. Glucosidases are plant cell-wall degrading enzymes shown to play a central role in fungal pathogenesis, as reported in various other fungi and oomycetes (Choi *et al.,* 2013). (6) *MGG_07444*, a multidrug resistance protein (MDR). The MDR transporters CgTpo1_1 and CgTpo1_2 have been shown to play a role in virulence of *Candida glabrata* (Santos *et al.,* 2016) (7) *MGG_15185*, Agglutinin isolectin 1 was found in four of the six analyses. Lectins mediate the attachment and binding of bacteria and viruses to their hosts. Similarly, it has been reported that fungal lectins can participate in the early stages of infection in humans (Houser *et al.,* 2013). (8) *MGG_05589*, a Lovastatin nonaketide synthase, which synthesize Lovastatin, a polyketide metabolite in *Aspergillus terreus*. Polyketides are major fungal secondary metabolites with varied biological activities, including being implicated in pathogenicity in plants (Collemare *et al.,* 2008). (9) *MGG_04289*, Dynamin GTPase, such as Mo*Dnm1* interact with partner proteins in the cytoskeleton and play important roles in appressorium function and pathogenicity in *M. oryzae* (Zhong *et al.,* 2016). Interestingly, none of these nine genes was found in the GWA analysis of the African cultivar NERICA5.

Six other genes (Table 4) were identified in the current study that have been either indirectly implicated in pathogenicity, or have not yet been fully functionally characterized to be involved in pathogenesis. (1) *MGG_04629*, an integral membrane protein was found across 4 of the 5 GWA analyses (2) *MGG_1497*1, an EF3 elongation factor was predicted in four of the six analyses. EF3’s are necessary for growth and development of an organism and thus essential for causing disease in the pathogenic fungus *C. albicans* (Perfect, 1996) (3) *MGG_00473,* rRNA-processing protein FCF2 has been identified in five of the six GWA analyses (4) Ubiquitin carboxyl-terminal hydrolase *MGG_05996*, a ubiquitin pathway gene that was found in three of the six analyses (5) *MGG_04954*, a stromal membrane associated protein was identified in 3 of 5 analyses (6) *MGG_14517,* the developmental regulator *flbA* was found in 3 of the 5 analyses.

This GWA study also identified multiple SNPs in predicted genes/hypothetical proteins/loci/repeat regions. The number of SNPs ranged from a maximum of nine to a minimum of two SNPs in any gene/genomic region (Table 5). Some of these genes repeatedly occurred in multiple studies. For example, *MGG_10232* had the presence of nine SNPs and was identified in three GWA analyses (IRBLTA CP1, IRBL3 CP4 and IRBL 9-W).

This study therefore demonstrates the power of using GWAS to identify markers of virulence in natural populations. This can provide insight into novel gene functions associated with rice blast disease that would otherwise not be identified based on conventional experimental analysis. In this way, it may be possible to identify key pathogenicity loci, or genes associated with overcoming resistance in cultivars being grown in SSA specifically. Further work will be necessary to test the roles of the identified genes in virulence on these cultivars and then to determine how resistance is conditioned in cultivars that are not susceptible.

## ACKNOWLEDGEMENTS

This project was supported, in part by the Sustainable Crop Production Research for International Development initiative, grant BB/J012157/1, funded jointly by the Biotechnology and Biological Sciences Research Council, UK, the Department for International Development and the Bill & Melinda Gates Foundation with additional funding from the Department of Biotechnology of India’s Ministry of Science and Technology. We thank the technical staff from T. Mitchell, J. C. Correll, and N. Talbot for their support.

## SUPPLEMENTARY DATA CAPTIONS

**Supplementary Table S1**: List of Genes/Genomic region homologs that encompass the significantly associated SNPs (500bp upstream and downstream of SNP locations) in the GWA analysis of NERICA5

**Supplementary Table S2**: List of Genes/Genomic region homologs that encompass the significantly associated SNPs (500bp upstream and downstream of SNP locations) in the GWA analysis of IRBLA-a

**Supplementary Table S3**: List of Genes/Genomic region homologs that encompass the significantly associated SNPs (500bp upstream and downstream of SNP locations) in the GWA analysis of IRBLTA-CP1

**Supplementary Table S4**: List of Genes/Genomic region homologs that encompass the significantly associated SNPs (500bp upstream and downstream of SNP locations) in the GWA analysis of IRBL3-CP4

**Supplementary Table S5**: List of Genes/Genomic region homologs that encompass the significantly associated SNPs (500bp upstream and downstream of SNP locations) in the GWA analysis of IRBLK-KA

**Supplementary Table S6**: List of Genes/Genomic region homologs that encompass the significantly associated SNPs (500bp upstream and downstream of SNP locations) in the GWA analysis of IRBL-9W

## REFERENCES

Altschul, S.F., Gish, W., Miller, W., Myers, E.W. & Lipman, D.J. (1990) “Basic local alignment search tool.” J. Mol. Biol. 215:403–410.

Baidya, S., Cary, J. W., Grayburn, W. S., Calvo A. M. (2011) Role of nitric oxide and flavohemoglobin homolog genes in Aspergillus nidulans sexual development and mycotoxin production. Appl. Environ. Microbiol. 77 5524–5528. 10.1128/AEM.00638-11.

Balasubramanian, V., Sie, M., Hijmans, R. J., & Otsuka, K. (2007) Increasing Rice Production in Sub-Saharan Africa: Challenges and Opportunities. Advances in Agronomy; Vol. 94, pp. 55–133 doi: 10.1016/S0065-2113(06)94002-4.

Bartoli, C, Roux, F. (2017) Genome-wide association studies in plant pathosystems: toward an ecological genomics approach. Front Plant Sci. 8:763.

Bradbury, P. J., Zhang, Z., Kroon, D. E., Casstevens, T. M., Ramdoss, Y., and Buckler, E. S. (2007) TASSEL: software for association mapping of complex traits in diverse samples. Bioinformatics 23, 2633–2635. doi: 10.1093/bioinformatics/btm308

Burdon, J.J., Thrall, P.H. (2009) Coevolution of plants and their pathogens in natural habitats. Science. 324:755–756.

Choi, J., Kim, K-T., Jeon, J. and Lee, Y-H. (2013) Fungal plant cell wall-degrading enzymedatabase: a platform for comparative andevolutionary genomics in fungi and Oomycetes. BMC Genomics; 14 (Suppl 5):S7.

Collemare, J., Billard, A., Böhnert, H.U., Lebrun, M.H. (2008) Biosynthesis of secondary metabolites in the rice blast fungus Magnaporthe grisea: the role of hybrid PKS-NRPS in pathogenicity. Mycol Res 112: 207–215.

Dalman, K., Himmelstrand, K., Olson, A., Lind, M., Brandstrom-Durling, M., Stenlid, J. (2013) A genome-wide association study identifies genomic regions for virulence in the non-model organism Heterobasidion annosum s.s. PLoS ONE 8:e53525. 10.1371/journal.pone.0053525.

Dean R.A. Talbot N.J. et al. (2005) The genome sequence of the rice blast fungus Magnaporthe grisea. Nature 434: 980–986.

Dodds P.N., Rafiqi M, Gan P.H.P, Hardham A.R, Jones D.A, Ellis J.G. (2009) Effectors of biotrophic fungi and oomycetes - pathogenicity factors and triggers of host resistance. New Phytology; 183:993–1000.

Dodds, P.N., and Rathjen, J.P. (2010) Plant immunity: towards an integrated view of plant–pathogen interactions. Nat Rev Genet 11: 539–548.

Elshire, R.J., Glaubitz, J.C., Sun, Q., Poland, J.A., Kawamoto, K., et al. (2011) A Robust, Simple Genotyping-by-Sequencing (GBS) Approach for High Diversity Species. PLoS One 6: e19379.

Fernández-Mazuecos, M., Mellers, G., Vigalondo, B., Sáez, L., Vargas, P., Glover, B. J. (2018) Resolving Recent Plant Radiations: Power and Robustness of Genotyping-by-Sequencing, Systematic Biology, Volume 67, Issue 2, Pages 250–268.

Gao, Y., Liu, Z., Faris, J. D., Richards, J., Brueggeman, R. S., Li X., et al. (2016) Validation of genome-wide association studies as a tool to identify virulence factors in Parastagonospora nodorum. Phytopathology 106, 1177–1185. 10.1094/PHYTO-02-16-0113-FI.

Gibriel, H.A., Thomma, B.P. and Seidl, M.F. (2016) The age of effectors: genome-based discovery and applications. Phytopathology, 106, 1206–1212.

Glaubitz J.C., Casstevens T.M., Lu, F., Harriman, J., Elshire, R.J., Sun, Q., et al. (2014) TASSEL-GBS: A High Capacity Genotyping by Sequencing Analysis Pipeline. PLoS ONE 9(2): e90346.

Guttman, D. S., McHardy, A. C., Schulze-Lefert, P. (2014) Microbial genome-enabled insights into plant-microorganism interactions. Nat. Rev. Genet. 15, 797–813. 10.1038/nrg3748.

Hartmann, F. E., Sanchez-Vallet, A., McDonald, B., Croll, D. (2017) A fungal wheat pathogen evolved host specialization by extensive chromosomal rearrangements. ISME J. 11, 1189–1204. 10.1038/ismej.2016.196.

Houser, J., Komarek, J., Kostlanova, N., Cioci, G., Varrot, A., Kerr, S.C., Lahmann, M., Balloy, V., Fahy, J.V., Chignard, M., et al. (2013) A soluble fucose-specific lectin from Aspergillus fumigatus conidia--structure, specificity and possible role in fungal pathogenicity. PLoS One. 8:e83077.

Hussain, W., Baenziger, P.S., Belamkar, V., Guttieri, M.J., Venegas, J.P., Easterly, A., Sallam, A., and Poland, J. (2017) Genotyping-by-Sequencing Derived High-Density Linkage Map and its Application to QTL Mapping of Flag Leaf Traits in Bread Wheat. Scientific Reports volume 7, Article number: 16394.

Jones, J. D., and Dangl, J. L. (2006) The plant immune system. Nature 444, 323–329.

Kamoun, S. (2007) Groovy times: Filamentous pathogen effectors revealed. Curr. Opin. Plant Biol. 10: 358–365.

Leboldus J. M., Kinzer K., Richards J., Ya Z., Yan C., Friesen T. L., et al. (2015) Genotype-by-sequencing of the plant-pathogenic fungi Pyrenophora teres and Sphaerulina musiva utilizing Ion Torrent sequence technology. Mol. Plant Pathol. 16 623–632. 10.1111/mpp.12214.

Lee, K.-T., So, Y.-S., Yang, D.-H., Jung, K.-W., Choi, J., Lee, D.-G., et al. (2016) Systematic functional analysis of kinases in the fungal pathogen Cryptococcus neoformans. Nature Communications, 7, 12766. http://doi.org/10.1038/ncomms12766.

Li, G., Huang, J., Yang, J., He, D., Wang, C., Qi, X., Taylor, I. A., Liu, J., and Peng, Y-L. (2017) Structure based function-annotation of hypothetical protein MGG_01005 from Magnaporthe oryzae reveals it is the dynein light chain orthologue of dynlt1/3. Scientific Reports volume 8, Article number: 3952.

Mgonja, E.M., Park, C.H., Kang, H, Balimponya, E.G., Opiyo, S.O., Bellizzi, M, Mutiga, S.K, Rotich, F., Devi Ganeshan, V., Mabagala, R., Sneller, C, Correll, J.C., Zhou, B., Talbot, N.J., Mitchell, T.K., and Wang, G-L. (2017) Genotyping-By-Sequencing-Based Genetic Analysis of African Rice Cultivars and Association Mapping of Blast Resistance Genes Against Magnaporthe oryzae Populations in Africa. Phytopathology. 107: 1039–1046. doi: 10.1094/PHYTO-12-16-0421-R.

Milgroom, M.G, Jiménez-Gasco, M.M, Olivares-García, C., Drott, M.T., Jiménez-Díaz, R.M. (2014) Recombination between clonal lineages of the asexual fungus Verticillium dahliae detected by genotyping by sequencing. PLoS ONE 9, e106740.

Monteil, C. L., Yahara, K., Studholme, D. J., Mageiros, L., Méric, G., Swingle, B., et al. (2016) Population-genomic insights into emergence, crop adaptation and dissemination of Pseudomonas syringae pathogens. Microbial Genomics, 2(10), e000089. http://doi.org/10.1099/mgen.0.000089. 28

Mutiga, S. K., Rotich, F., Devi Ganeshan, V., Mwongera, D. T., Mgonja, E. M., Were, V. M., Harvey, J. W., Zhou, B., Wasilwa, L., Feng, C., Ouédraogo, I., Wang, G.- L., Mitchell, T. K., Talbot, N. J., and Correll, J. C. (2017) Assessment of the Virulence Spectrum and Its Association with Genetic Diversity in Magnaporthe oryzae Populations from Sub-Saharan Africa. Phytopathology 107: 7, 852-863.

Nalley, L., Tsiboe, F., Durand-Morat, A., Shew, A., Thoma, G. (2016) Economic and Environmental Impact of Rice Blast Pathogen (Magnaporthe oryzae) Alleviation in the United States. PLOS ONE 11(12): e0167295. https://doi.org/10.1371/journal.pone.0167295.

Nigatu, G., Hansen, J., Childs, N., and Seeley, R. (2017) Sub-Saharan Africa Is Projected To Be the Leader in Global Rice Imports. Amber Waves.

Norelli, J. L., Wisniewski, M., Fazio, G., Burchard, E., Gutierrez, B., Levin, E., and Droby, S. (2017) Genotyping-by-sequencing markers facilitate the identification of quantitative trait loci controlling resistance to Penicillium expansum in Malus sieversii. PLoS ONE, 12(3), e0172949. http://doi.org/10.1371/journal.pone.0172949.

Occhipinti, A. (2013) Plant coevolution: evidences and new challenges. J Plant Interact 8: 188–196.

Patterson., N, Price., AL, Reich., D. (2006) Population Structure and Eigenanalysis. PLoS Genet 2(12): e190.

Pearce, L. R., Komander, D. and Alessi, D. R. (2010) The nuts and bolts of AGC protein kinases. Nat. Rev. Mol. Cell Biol. 11, 9–22.

Perfect, J.R. (1996) Fungal virulence genes as targets for antifungal chemotherapy. Antimicrob. Agents Chemother. 40, 1577 1583.

Petit-Houdenot, Y. and Fudal, I. (2017) Complex interactions between fungal avirulence genes and their corresponding plant resistance genes and consequences for disease resistance management. Front. Plant Sci. 8, 1072. doi: 10.3389/ fpls.2017.01072.

Pilet-Nayel, M-L., Moury, B., Caffier, V., Montarry, J., Kerlan, M-C., Fournet, S., Durel, C-E., Delourme, R. (2017) Quantitative resistance to pathogens in plant pyramiding strategies for durable crop protection. Front. Plant Sci. 8:1838.

Plissonneau, C., Blaise, F., Ollivier, B., Leflon, M., Carpezat, J., Rouxel, T., et al. (2017) Unusual evolutionary mechanisms to escape effector-triggered-immunity in the fungal phytopathogen Leptosphaeria maculans. Mol. Ecol. 26, 2183–2198. 10.1111/mec.14046.

Quoc, N. B. and Chau, N. N. B. (2017) The role of cell wall degrading enzymes in Pathogenesis of Magnaporthe oryzae. Current Protein and Peptide Science, 18, 1–16.

R Core Team (2017) R: A language and environment for statistical computing. Vienna, Austria: R Foundation for statistical computing 2016; 860–864.

Rafiei, V., Banihashemi, Z., Jiménez-Díaz, R. M., Navas-Cortés, J. A., Landa, B. B., Jiménez-Gasco, M. M., Turgeon, B. G., Milgroom, M. G. (2017) Comparison of genotyping by sequencing and microsatellite markers for unravelling population structure in the clonal fungus Verticillium dahlia. Plant Pathology 67, 76–86.

Ray, S., Singh, P. K., Gupta, D. K., Mahato, A. K., Sarkar, C., Rathour, R., Singh, N. K., Sharma, T. R. (2016) Analysis of Magnaporthe oryzae genome reveals a fungal effector, which is able to induce resistance response in transgenic rice line containing resistance gene, pi54. Frontiers in Plant Science, 7, p. 1140.

Rempola, B., Karkusiewicz, I., Piekarska, I., Rytka, J. (2006) Fcf1p and Fcf2p are novel nucleolar Saccharomyces cerevisiae proteins involved in pre-rRNA processing. Biochem. Biophys. Res. Commun.; 346:546–554.

Saito, K., Nelson, A., Zwart, S. J., Niang, A., Sow, A., Yoshida, H., and Wopereis, M. C. S. (2013) Towards a Better Understanding of biophysical determinants of yield gaps and the potential for expansion of the rice area in Africa. Realizing Africa’s Rice Promise. M. C. S. Pages 188–203.

Samalova, M., Johnson, J., Illes, M., Kelly, S., Fricker, M., and Gurr, S. (2013) Nitric oxide generated by the rice blast fungus Magnaporthe oryzae drives plant infection. New Phytol. 197, 207–222. doi: 10.1111/j.1469-8137.2012.04368.x

Sanchez-Vallet, A., Hartmann, F.E., Marcel T. C., and Croll, D. (2018) Nature’s genetic screens: using genome-wide association studies for effector discovery. Molecular Plant Pathology, 19 (1): 3-6, Oxford: Wiley-Blackwell, 2018.

Santos R., Costa C., Mil-Homens D., Romão D., de Carvalho C. C. C. R., Pais P., et al. (2017) The multidrug resistance transporters CgTpo1_1 and CgTpo1_2 play a role in virulence and biofilm formation in the human pathogen Candida glabrata. Cell. Microbiol. 19, 1–13. 10.1111/cmi.12686.

Seifbarghi, S., Borhan, M.H., Wei, Y., Coutu, C., Robinson, S.J., and Hegedus, D.D. (2017) Changes in the Sclerotinia sclerotiorum transcriptome during infection of Brassica napus. BMC Genomics. Mar 29; 18 (1):266.

Sharma, T. R., Rai, A. K., Gupta, S. K., Vijayan, J., Devanna, B. N., Ray, S. (2012). Rice blast management through host-plant resistance: retrospect and prospects. Agri. Res. 1, 37–52. 10.1007/s40003-011-0003-5.

Talas, F., Kalih, R., Miedaner, T., McDonald, B. A. (2016) Genome-wide association study identifies novel candidate genes for aggressiveness, deoxynivalenol production, and azole sensitivity in natural field populations of Fusarium graminearum. Mol. Plant Microbe Interact. 29, 417–430.

Tharreau, D., Fudal, I., Andriantsimialona, D., Santoso, Utami, D., Fournier, E., Lebrun, M.H., Nottéghem, J.L. (2009) World population structure and migration of the rice blast fungus, Magnaporthe oryzae. Advances in genetics, genomics and control of rice blast disease. Wang GL, Valent B, eds. Dordrecht, the Netherlands: Springer, 209–215.

Torkamaneh, D., Laroche, J., Bastien, M., Abed, A. and Belzile, F. (2017a) Fast-GBS: a new pipeline for the efficient and highly accurate calling of SNPs from genotyping-by-sequencing data. BMC Bioinformatics, 18, 5.

Wang, Q., Chen, D., Wu, M., Zhu, J., Jiang, C., Xu, J-R and Liu, H. (2018) MFS Transporters and GABA Metabolism Are Involved in the Self-Defense Against DON in Fusarium graminearum. Front. Plant Sci., 13 April 2018.

Wu, J.Q., Sakthikumar, S., Dong, C., Zhang, P., Cuomo, C.A. and Park, R.F. (2017) Comparative genomics integrated with association analysis identifies candidate effector genes corresponding to Lr20 in phenotype-paired Puccinia triticina isolates from Australia. Front. Plant Sci. 8, 148.

Wu, Y., Xiao, N., Yu, L., Pan, C., Li, Y., Zhang, X., et al. (2015) Combination patterns of major R genes determine the level of resistance to the M. oryzae in rice (Oryza sativa L.). PLoS ONE 10:e012613010.1371/journal.pone.0126130.

Zhong, K., Li, X., Le, X., Kong, X., Zhang, H., Zheng, X., Wang, P. and Zhang, Z. (2016) MoDnm1 Dynamin Mediating Peroxisomal and Mitochondrial Fission in Complex with MoFis1 and MoMdv1 Is Important for Development of Functional Appressorium in Magnaporthe oryzae. PLoS Pathog 12(8): e1005823. https://doi.org/10.1371/journal.ppat.1005823.

